# Short template switch events explain mutation clusters in the human genome

**DOI:** 10.1101/038380

**Authors:** Ari Löytynoja, Nick Goldman

**Affiliations:** Institute of Biotechnology, University of Helsinki, Helsinki, Finland; European Molecular Biology Laboratory, European Bioinformatics Institute (EMBL-EBI), Wellcome Genome Campus, Hinxton, UK

**Keywords:** *de novo* assembly, hairpin loop structures, human evolution, human resequencing, multi-nucleotide mutations, mutation clusters, template switch events

## Abstract

Resequencing efforts are uncovering the extent of genetic variation in humans and provide data to study the evolutionary processes shaping our genome. One recurring puzzle in both intra- and inter-species studies is the high frequency of complex mutations comprising multiple nearby base substitutions or insertion-deletions. We devised a generalized mutation model of template switching during replication that extends existing models of genome rearrangement, and used this to study the role of template switch events in the origin of such mutation clusters. Applied to the human genome, our model detects thousands of template switch events during the evolution of human and chimp from their common ancestor, and hundreds of events between two independently sequenced human genomes. While many of these are consistent with the template switch mechanism previously proposed for bacteria but not thought significant in higher organisms, our model also identifies new types of mutations that create short inversions, some flanked by paired inverted repeats. The local template switch process can create numerous complex mutation patterns, including hairpin loop structures, and explains multi-nucleotide mutations and compensatory substitutions without invoking positive selection, complicated and speculative mechanisms, or implausible coincidence. Clustered sequence differences are challenging for mapping and variant calling methods, and we show that detection of mutation clusters with current resequencing methodologies is difficult and many erroneous variant annotations exist in human reference data. Template switch events such as those we have uncovered may have been neglected as an explanation for complex mutations because of biases in commonly used analyses. Incorporation of our model into reference-based analysis pipelines and comparisons of *de novo*-assembled genomes will lead to improved understanding of genome variation and evolution.

## Introduction

Mutations are not evenly distributed in genome sequences. Base substitutions and short insertions and deletions (‘indels’, up to tens of bp in length) usually reflect errors in DNA replication and/or repair (Gu et al., 2008) and tend to form clusters (e.g. Averof et al., 2000; Whelan and Goldman, 2004; Harris and Nielsen, 2014; Sudmant et al., 2015). Explanations for these monogenic point mutation clusters (subsequently referred to as simply ‘mutation clusters’) vary from an error-prone polymerase (Harris and Nielsen, 2014) to indels being mutagenic (Tian et al., 2008).

Genomic rearrangements are defined as gross DNA changes, typically thousands to millions of base pairs and covering multiple different genes (Gu et al., 2008). Although difficult to study using traditional genome sequencing methods, they have recently become the focus of intense research (e.g. Pendleton et al., 2015; Sudmant et al., 2015) due to the advent of next generation sequencing techniques and the importance of their effects in both somatic and germ cells, causing cancers and genetic diseases. Earlier mechanisms proposed to explain genomic rearrangements were ones involving recombination, in particular non-allelic homologous recombination (NAHR) and non-homologous end-joining (NHEJ; see reviews by Gu et al., 2008, and Carvalho and Lupski, 2016). More recently, replication-based mechanisms such as serial replication slippage (SRS; Chen et al., 2005*c*,*a*,*b*), break-induced replication (BIR; Morrow et al., 1997), fork stalling and template switching (FoSTeS; Lee et al., 2007) and microhomology-mediated break-induced replication (MMBIR; Hastings, Ira and Lupski, 2009) have been proposed. Mutations attributed to all of these mechanisms typically involve major genomic rearrangements (see also Costantino et al., 2013, and Carvalho and Lupski, 2016).

Gu et al. (2008) and Carvalho and Lupski (2016) suggest that replication-based genome rearrangement mechanisms could be responsible for both small-scale and large-scale mutations, and that their implications for evolution have yet to be investigated. We hypothesized that a generalized model of genome mutation that encompassed the consequences of replication-based mechanisms such as SRS, BIR, FoSTeS and MMBIR might be able to account for the observation of mutation clusters in higher organisms more parsimoniously than invoking a process of successive base substitutions and indels in a small region.

Common to all these mechanisms is that during replication the 3' end of the nascent DNA strand dissociates from the original template and invades another (physically close) open replication fork. A segment is incorporated using this new template until the strand dissociates again. Replication may continue through a complex series of such template ‘switch-and-return’ events; eventually the nascent DNA reassociates with the original template and replication proceeds as normal. Complex examples with multiple switches have been convincingly demonstrated by (e.g.) Chen et al. (2005*a*) and Lee et al. (2007).

Long-range template switch events are those that where the nascent DNA changes template between distinct replication forks and the inserted segment(s) derive from genomic regions thousands to millions of bp distant (Lee et al., 2007) or even from other chromosomes (Chen et al., 2005*c*,*a*; see also Smith et al., 2007). These have been linked with large genomic rearrangements under mechanisms such as BIR, FoSTeS and MMBIR. Short-range events involve template switches within the same replication fork, meaning the inserted segments derive from regions nearby in primary sequence. These have been considered previously in a limited manner as a possible explanation of mutation clusters. In bacteria, mutations creating perfect inverted repeats occur with high frequency (Dutra and Lovett, 2006) and are thought to involve intra-strand template switching, where the nascent strand is itself used as the template, or inter-strand template switching, where the strand complementary to the original template is used (Fig. 1a, b) (Ripley, 1990). Such template switching is believed to require a pre-existing near-perfect inverted repeat, which is converted into a perfect inverted repeat within the nascent strand by the use of complementary sequence for the transient template.

**Fig. 1:**
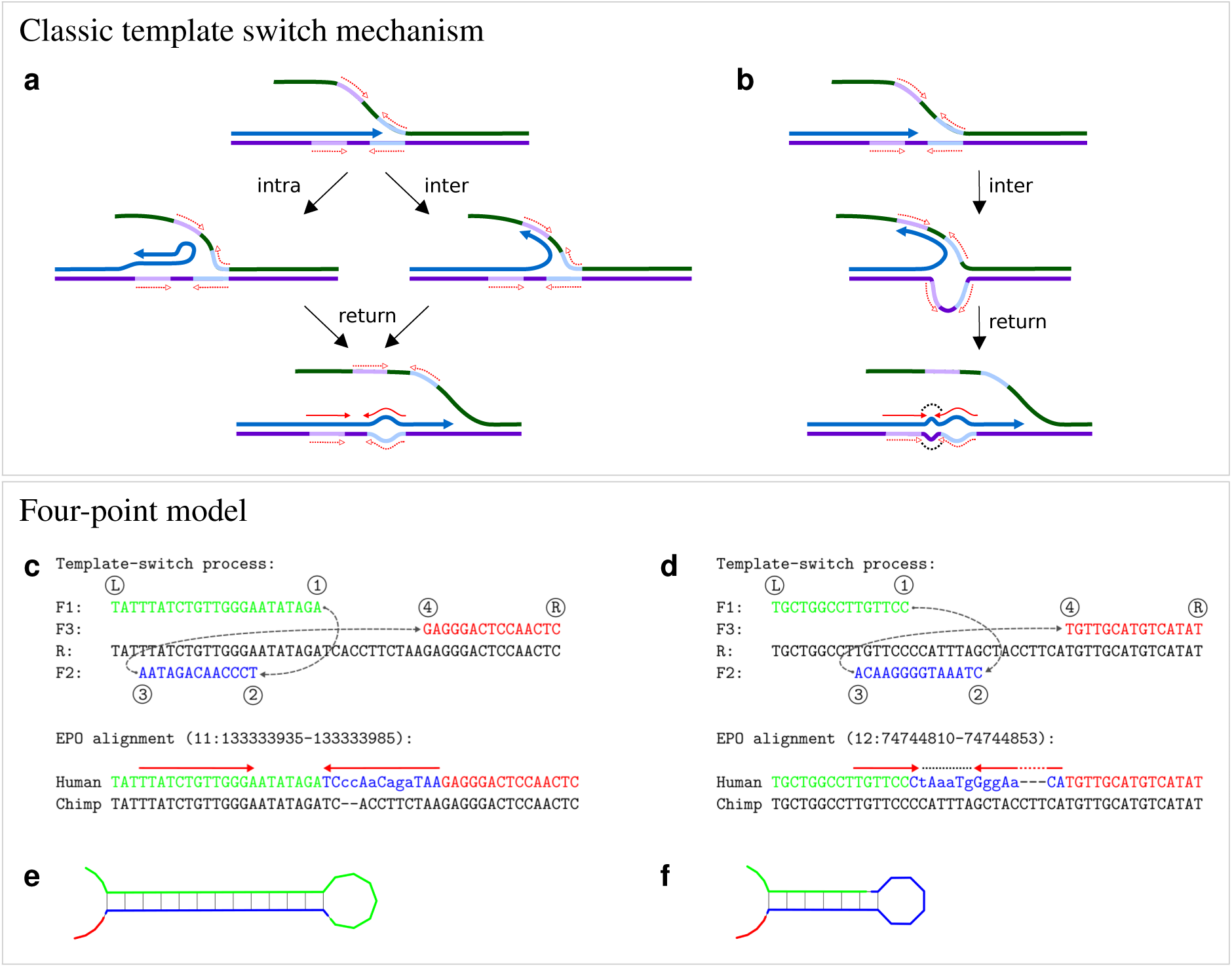
Classic template switch mechanism and the new four-point model. **a**, **b**, The classic template switch mechanism creates perfect inverted repeats. **a**, DNA replication (blue arrow) exchanges template and converts a nearly perfect inverted repeat (dashed red arrows) into a perfect one (solid red arrows), causing a cluster of differences (bulge, bottom); this can happen by an intra-strand (left) or an inter-strand (right) switch. **b**, An inter-strand switch may invert the spacer of the repeat (black dots). **c**, **d**, The new four-point model generalizes the template switch mutation process. Template exchanges are described with four switch points (labelled ①→② projected onto a reference sequence (R). The points define three sequence fragments (F1–F3) which, when concatenated, create a mutated output (mismatches shown in lower case in the human sequence). F1 and F3 are copied from R; F2 is copied complementary to either F1 (intra-strand switch) or R (inter-strand switch). The model can perfectly explain complex mutations observed in real data (bottom). **c**, Event “3-2-1-4", named for the order of the switch points along R, creates an inverted repeat (bottom; red arrows). **d**, Event “3-1-2-4" creates an inverted repeat (red arrows) separated by an inverted spacer (dotted line). **e**, **f**, Predicted secondary structures generated by the inverted repeats created in the Human sequences in **c**, **d**, respectively.

Links to original data:

**c**: http://grch37.ensembl.org/Homo%5Fsapiens/Location/Compara%5FAlignments?align=548&r=11:133333935-133333985

**d**: http://grch37.ensembl.org/Homo%5Fsapiens/Location/Compara%5FAlignments?align=548&r=12:74744810-74744853

Under this model, both intra- and inter-strand template switch types can cause sequence changes within the repeat (Fig. 1a), while the latter can additionally invert the ‘spacer’ sequence (the region between the repeat fragments; Fig. 1b). While these changes can appear as clusters of differences (Dutra and Lovett, 2006) and have been detected in genes implicated in human genetic disease (Chen et al., 2005*b*), this bacterial-style mechanism has not been considered significant in the evolution of higher organisms (Ladoukakis and Eyre-Walker, 2008). These conclusions were based on limited data, however, and on an assumption that the mechanism necessarily creates perfect inverted repeats. We compared human and chimp genomes and observed mutation clusters that create novel inverted repeats consistent with the bacterial mechanism. Many clusters could only partially be explained by the creation of an inverted repeat, however, and novel repeats were often flanked by indels or dissimilar sequence, inconsistent with the classical model.

Even with the underlying biological mechanism uncertain, we realized that the existence and properties of a template switch mutation process, capable of creating inverted repeats, could be studied using pairs of closely-related genome sequences. Specifically, we devised the ‘four-point model’ of template switching, based on short-range switch-and-return events. The model is computationally tractable for genome-wide searches, permitting the discovery of DNA within a short distance that can explain the existence of a mutation cluster as a single template switch event. Events detected by our model can be validated by controls that determine the background level of hits expected due to the possibility of local genomic regions by chance containing the sequence needed to match a mutation cluster.

We first apply our method to genome-wide alignments of human and chimp. Focusing on the complex and unique regions of the genome and comparing the solutions involving a template switch to the original linear sequence alignments, we find that thousands of mutation clusters can be explained as the result of a short-range template switch event during replication. Next, comparing two *de novo*-assembled human genomes, we detect nearly 270 mutation clusters that are candidates for template switch events, including numerous polymorphic loci. Finally, we investigate further evidence for these last mutations in variant calls from the 1000 Genomes (1kG) project. Many of them are indeed observed in the 1kG data; numerous inconsistencies are explained as artefacts of erroneous mapping and variant calling. This calls into question the accuracy of current reference-based mapping strategies for population resequencing and consequent inferences about local mutation rates. It highlights the need for highly accurate assemblies, using either *de novo* methods or improved mapping strategies based on a better understanding of the mutation processes acting on genomes.

## Results and discussion

### Four-point model of template switching

Any single template switch-and-return event can be described by a model that projects four sequence positions onto a reference sequence and then constructs a replication copy from the three fragments defined by these points. For convenience, we describe the process as involving the nascent leading strand; the model equally well describes events corresponding to the lagging strand. We have implemented the model with the assumption that template switches are short-range (i.e. use the same replication fork) and involve ‘jumps’ in the replication process to use a template strand other than the original one (‘replication slippage in *trans*’ in the terminology of Chen et al., 2005*b*). This can be the nascent DNA strand itself (intra-strand switching), or the lagging strand (inter-strand switching). We do not attempt to use the model to explain long-range template switches, or multiple successive rounds of template switching. While the four-point model could in principle be extended to cover these possibilities including all of the outcomes that may arise from the SRS, BIR, FoSTeS and MMBIR mechanisms, it would be computationally intractable, and unlikely to find compelling examples given that essentially the entire genome would be available as the possible explanation of a relatively small number of base substitutions and indels.

The four-point model is illustrated in Fig. 1c–d. Assuming that replication proceeds from left 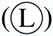 to right 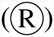, points ① and ② indicate the location of the first switch event with the nascent strand dissociating from the leading strand at location ① and continuing at ② lagging strand, or equivalent location on the nascent strand). Similarly, the second (return) switch event comprises a second dissociation taking place at ③ and reassociation with the leading strand at ④. The replication copy then consists of fragments 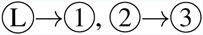 (complemented, note), 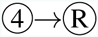 (Fig. 1c–d). If fragment ②→③ overlaps with fragment 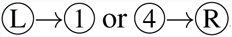, the mutation creates a novel inverted repeat that then may be capable of forming a RNA secondary structure (Fig. 1e–f).

Modeling the template switch process like this has two major advantages. First, it allows for a formal analysis of mutation events and their evaluation in comparison to alternative explanations. Second, our description of the process is general and has few *a priori* constraints for the template exchanges. Our projection of switch points onto a reference is impartial regarding the type of the switch event—either intra- or inter-strand—and the model only requires that the ②→③ fragment is copied in reverse-complement orientation. The possible outcomes under the four-point model are defined by the relative order and distance of the switch points, and the classical mechanism proposed to explain inverted repeats in bacteria is a special case of our generalized model (cf. Fig. 1a,c). Supplementary Fig. 1 illustrates all the possible cases under the model, covering the scenarios described before (Fig. 1a–b) as well as several others, including creation of inverted and direct repeats flanked by dissimilar sequence and one case causing inversion of a sequence fragment only. For creation of mutation clusters, an important characteristic of the model is that replacement of the ①→④ fragment with the reverse-complement of the ②→④ fragment by a single switch event can generate changes that, when viewed in a linear alignment, will appear as multiple nearby substitutions and indels.

### Application of the four-point model

To test whether biological data support the proposed model as an explanation of mutation clusters in the human genome, we implemented a computational tool based on a custom dynamic programming (dp) algorithm. The tool identifies clusters of differences between two aligned genomic sequences and then searches for an explanation of the region of dissimilarity in one sequence (replicate output) by copying a fragment from the other sequence (reference) in reverse-complement orientation, as achieved in the four-point model. With two closely-related sequences, parallel mutation will be rare and we arbitrarily designate one sequence as the reference and assume that it represents the ancestral form around each mutation event in the replicate lineage. For full details of the dp algorithm used to determine the optimal four-point model explanation for each mutation cluster, see Methods, Supplementary Fig. 2 and Supplementary Algorithm 1.

We focused on the complex and unique regions of human and chimp genomes and compared the solutions involving a template switch to the original linear sequence alignments. From the potential cases of template switch events detected, we filtered sets of high-confidence events (see Methods). To create a control to assess false positives, we used a proxy for observing the mutation patterns by chance: we computed the best solutions explaining the dissimilar sequence regions with the fragment ②→③ copied in reverse (i.e. not reverse-complement) orientation and evaluated these solutions using the same criteria. Modification of our dp algorithm to achieve these controls is also described in Supplementary Algorithm 1.

### Discovery of four-point mutation events from human-chimp data

We applied our model to genome-wide Ensembl EPO alignments (v.71, 6 primates) of human and chimp (Flicek et al., 2013; Paten et al., 2008), considering the chimp sequence the reference and the human sequence the mutated copy. The portion of human-chimp alignment data not masked as repeats or low-complexity sequence (48.5% of total length; see Methods) contains 14.51*×*10^6^ base differences and 1.19*×*10^6^ indels. Of these, 3.84*×*10^6^ base differences (26.4%) and 0.76 *×* 10^6^ indels (63.9%) are within mutation clusters consisting of multiple nearby base differences or alignment gaps. Using our computational tool, we found 4,778 candidate four-point mutation events, spread across all human chromosomes, overlapping with 11,723 base differences and 1,288 indels, or 0.31% and 0.17% of total clustered unmasked events, respectively. Some candidate events were consistent with the original mechanism proposed for bacteria and convert a near-perfect inverted repeat into a perfect one (see example in Fig. 2a–b) but the majority were associated with large sequence changes, causing multiple base differences and indels in linear alignments (e.g. Fig. 2c–d). While any complex mutation could be generated by a combination of simple, ‘traditional’, mutations, Occam’s razor suggests that a four-point model template switch mutation is a better explanation than multiple substitutions and indels occurring in such a cluster. However, we also noticed that matches shorter than 12– 13 bases are often found by chance (Supplementary Figs. 3, 4) and, despite strict filtering (see Methods), our list of candidate events might still contain false positives. To get an unbiased picture of the process, we removed events with ②→③ fragment shorter than 14 bases. This was done to improve the signal to noise ratio and does not mean that short template switch events could not happen: in contrast, many cases with a short ②→③ fragment appear highly convincing (e.g. Fig. 1c).

**Fig. 2:**
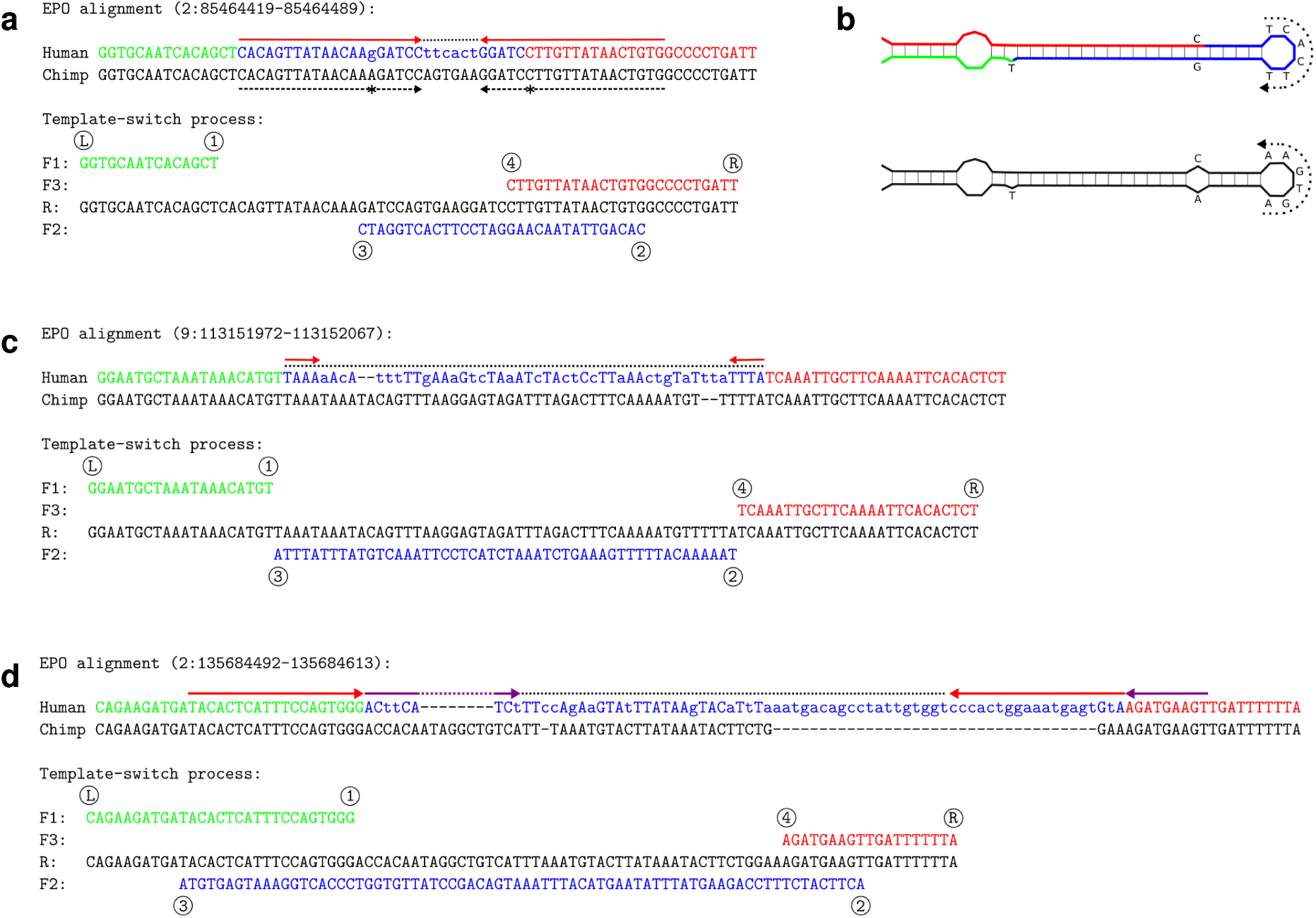
Example events detected in human. **a**, A near-perfect inverted repeat in chimp (dashed black arrows, the one mismatch indicated with asterisks) has been converted into a perfect inverted repeat (red arrows) in human (top). The cluster of six additional dissimilarities (dotted line) in fact represents perfect inversion of the 6-bp spacer sequence and makes the template switch (bottom) a likely explanation. **b**, Predicted DNA secondary structure before (chimp; bottom) and after (human; top) the template switch event. The dotted arrows indicate the reverse-complemented spacer region, which the four-point model explains with a single event. **c**, **d**, Additional complex mutation patterns (mismatches in lower case) that can be explained by a single template switch event. **c**, Event “1-3-2-4" only converts the spacer sequence. **d**, Event “3-1-4-2" converts the spacer sequence and creates two inverted repeats (red and magenta arrows).

Links to original data:

**a**: http://grch37.ensembl.org/Homo%5Fsapiens/Location/Compara%5FAlignments?align=548&r=2:85464419-85464489

**c**: http://grch37.ensembl.org/Homo%5Fsapiens/Location/Compara%5FAlignments?align=548&r=9:113151972-113152067

**d**: http://grch37.ensembl.org/Homo%5Fsapiens/Location/Compara%5FAlignments?align=548&r=2:135684492-135684613

After this filtering, we assigned the 794 remaining candidate events to specific event types based on the relative positions of the switch points and computed their frequencies. We found that, of the 12 possible conformations of switch points, only six are present (Table 1, human vs. chimp comparison). Of these, two event pairs are ‘mirror cases’ indistinguishable from one-another if both leading and lagging template strands are considered (see Supplementary Fig. 1), and the six conformations observed therefore define four distinct switch event types. Type “1-4-3-2” (with its mirror case “3-2-1-4”; Supplementary Fig. 1a; e.g. Fig. 1c) creates an inverted repeat and accounts for 31% of the high-confidence events detected in the chimp-human comparison. Type “1-3-4-2” (with its mirror case “3-1-2-4”; Supplementary Fig. 1a; e.g. Fig. 1d) creates an inverted repeat separated by an inverted spacer sequence, accounting for 23% of events. The remaining two types are novel and only achievable under our four-point model: type “1-3-2-4”, accounting for 45% of events, only inverts a sequence fragment and creates no repeat (e.g. Fig. 2c), and type “3-1-4-2” creates two inverted repeats separated by an inverted spacer (e.g. Fig. 2d) and accounts for 1% of events.

**Table 1:** **Proportion of event types.** Proportion of different event types among the high-confidence cases, for the comparisons of human vs. chimp and of two humans. Only one observed event type could happen via intra-strand switching (red star, its mirror case indicated with a black star). All other events can only happen inter-strand (see also Supplementary Fig. 1).

| event type | output | human vs.<br>chimp | two<br>humans |
| --- | --- | --- | --- |
| ★ 1-4-3-2, ★ 3-2-1-4 | inverted repeat | 0.31 | 0.37 |
| 1-3-4-2, 3-1-2-4 | inverted repeat and inverted spacer | 0.23 | 0.15 |
| 1-3-2-4 | inverted fragment | 0.45 | 0.47 |
| 3-1-4-2 | two inverted repeats and inverted spacer | 0.01 | 0.01 |
| events total |  | 794 | 92 |

The unifying feature of the event types theoretically possible under the model but not observed in real sequence data is that in the ordering of the switch points, ④ precedes ①. This is the hallmark of an event in which the second (return) template switch requires the opening of the newly synthesized DNA double helix (see Supplementary Fig. 1). In addition we observe numerous cases of inversion of spacer sequences; this cannot occur when ② precedes ①, a prerequisite of intra-strand switches. These discoveries suggest that template switches occur inter-strand: that is, the fragment ②→③ is copied from the opposite strand (Fig. 1).

Although inversions of spacer sequences have been observed in bacteria (Ripley, 1990), the intra-strand mechanism has been the dominant hypothesis (Dutra and Lovett, 2006). It appears that this is not correct, at least for evolution since the human-chimp divergence. We also find that the relative frequencies of different event types are very different. In part this may be determined by factors such as the length distribution of the copied fragment (Supplementary Fig. 3) and type “3-1-4-2” requiring that the fragment ②→③ overlaps with both ① and ④. However, the frequencies of different event types may also reflect the properties of the mutation process, e.g. template switching benefiting from the proximity of the DNA strands, or the chance of the new mutation escaping error correction.

### Identification of polymorphic mutations in human data

To understand whether template switch events are actively shaping human genomes, we analyzed human resequencing data and searched for polymorphic loci. We first aligned the human reference genome (GRCh37) to that of a Caucasian male (Venter, also denoted HuRef; Levy et al., 2007), both based on classical capillary sequencing and assembled independently. We then considered Venter as the reference and identified clusters of mutations in GRCh37 that were consistent with different types of four-point model template switch events. Using the same approach as in the human-chimp comparisons, we identified 267 candidate events in the unmasked portion of the human genome and then selected a smaller set of high-confidence cases for a more close analysis (see Methods). For these 92 events, the proportions of different event types were similar to those found in human-chimp comparisons. Again, only the six types not requiring opening of the new helix were found and the majority of events require inter-strand switches (Table 1: two humans comparison).

Still focusing on these 92 candidate events, we manually studied the Caucasian male sequence data mapped onto the reference genome (Li and Durbin, 2011). We could resolve the genotype of the Caucasian male for 76 (83%) of the candidate events and found 40 of them heterozygous, i.e. the sequence data contain reads consistent with both Venter and GRCh37 alleles (Fig. 3a, b; Supplementary Table 1; see also Supplementary Data 1 available at http://loytynojalab.biocenter.helsinki.fi/software/fpa). In two cases the read data revealed that the mutations forming the cluster are not linked and are the result of two independent mutation events (Supplementary Fig. 5) and in the remaining 16 cases, mapping of Venter sequence reads against GRCh37 was inconsistent with the alignment of genome assemblies; we did not consider these ambiguous mutations any further.

**Fig. 3:**
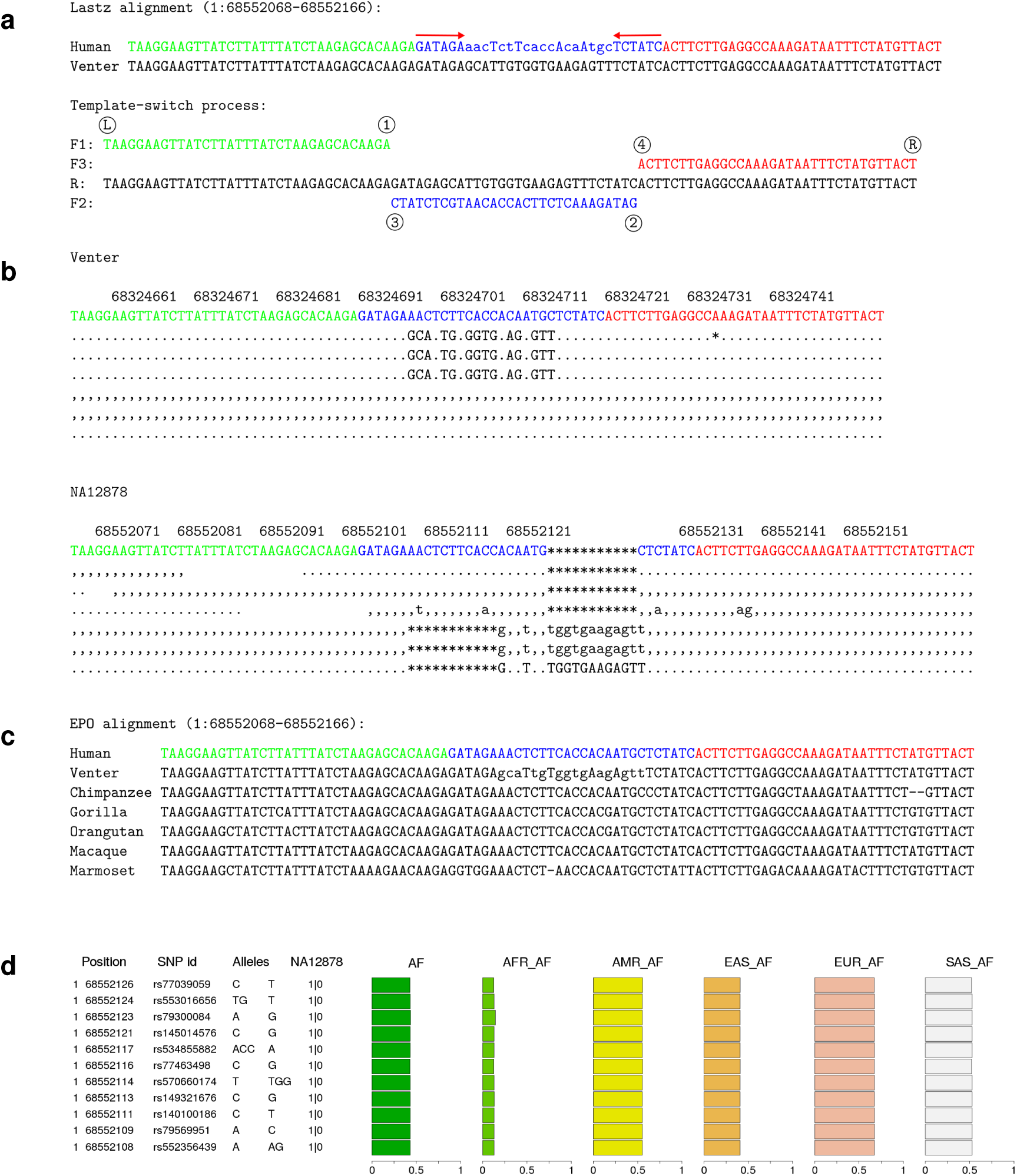
A template switch mutation event with variable allele frequencies in human populations. **a**, Four-point model explanation of a complex mutation between the human reference GRCh37 (denoted Human) and a Caucasian male (Venter). Notation is as in Fig. 1. **b**, A subset of the original sequencing reads from the Caucasian male (top) and the 1kG individual NA12878 (bottom). Dots and commas indicate the read matching to the reference on the forward and reverse strand, upper- and lower-case characters denote the corresponding mismatches, and asterisks mark the alignment gaps. These reads reveal heterozygosity at the locus. **c**, The EPO alignment for primates reveals that the human reference (Human) is the ancestral form. As all other primates resemble the reference allele, the most parsimonious explanation is that the mutation (Venter) has happened in the human lineage since its divergence from the human-chimp ancestor. **d**, 1kG variation data explain this event as a cluster of 7 single nucleotide polymorphisms and 4 indels. The phased genotypes for NA12878 (1 0) indicate that the variant alleles are linked and all in the same haplotype. The single origin of the whole cluster is further supported by the uniform derived allele frequencies across the sites within all 1kG-data (AF) and within each superpopulation (AFR, AMR, EAS, EUR, SAS).

We then looked at the same loci in the 1kG data (1000 Genomes Project Consortium et al., 2015) and studied the alignment data for individual NA12878. We found that NA12878 has a non-reference allele at 47 of the 76 resolved loci (62%) and, with the exception of the two cases mentioned, all the changes are found within the same sequence reads. This finding has two implications. First, with two different sequencing technologies (capillary and Illumina) and analysis pipelines showing the same mutation patterns, we can reject the possibility that the observed events could be technical artefacts. Second, the agreement of short-read data with assembly based on long capillary reads suggests that the template switch mutation model can be studied using modern resequencing data.

### Elimination of mutation accumulation hypothesis

In principle, the perfect linkage of adjacent sequence changes in two unrelated individuals could also be explained by mutations being accumulated over a long period of time in complete absence of recombination. To rule that out, we assessed the maximum age of the mutation clusters using phylogenetic information (Fig. 3c). The EPO alignments contain data from at least two additional primate species for 73 loci. The two alleles detected between the two humans GRCh37 and Venter segregate among the primate species in only one of these loci; in all 72 other cases, all primate sequences resemble one of the two human alleles while the second human allele is unique (Fig. 3c; Supplementary Data 2, http://loytynojalab.biocenter.helsinki.fi/software/fpa). Although some loci could be polymorphic in non-human primates, the result suggests that a great majority of the events are young and the adjacent changes within the mutation clusters result from single mutation events.

### Mutation clusters in 1000 Genomes variation data

NA12878 is only one individual and a greater proportion of the 76 candidate loci may be truly polymorphic in larger samples. We investigated whether the mutations caused by template switch events are visible in variation data. Using the 1kG variant calls (1000 Genomes Project Consortium et al., 2015) we found that this is indeed the case: of the 76 confirmed events between the reference and the Caucasian male, the mutation pattern created by the event is completely explained by combinations of the 1kG variants (separate calls of indels and single nucleotide polymorphisms) at 35 loci, and partially explained at a further 16 loci. In most cases, the mutations at a locus have uniform allele frequencies within human populations, further demonstrating the perfect linkage and the single origin for the full mutation cluster (Fig. 3d; Supplementary Data 1, http://loytynojalab.biocenter.helsinki.fi/software/fpa). The variation data confirm the two earlier cases as combinations of independent mutations (Supplementary Fig. 5) but, for all other inconsistencies, alignment data show the incomplete mutation patterns and the non-uniform allele frequencies to be artefacts from erroneous mapping and variant calling (Supplementary Fig. 6). Such inconsistencies may be expected when the variant calls are based on mapping of short reads containing multiple differences to a reference sequence, and demonstrate the difficulty of correctly detecting complex mutations using current reference-based analysis methods.

Despite highly uniform allele frequencies, the 1kG variant calls consider the template switch events that we identified to be clusters of independent mutations events—the largest clusters consisting of more than ten apparently independent mutation events (e.g. Supplementary Fig. 7)—and thus seriously exaggerate the estimates of local mutation rate. On the other hand, uniform allele frequencies at adjacent positions indicate a shared history for a mutation cluster and potentially allow computational detection of events. To test this, we investigated whether any of the events found between human and chimp are still polymorphic in humans and associated with a cluster of SNP positions with uniform allele frequencies (see Methods). We found several such events, the frequencies of the two haplotypes varying from close to 0 to nearly 1, and the frequencies differing significantly between populations (Supplementary Fig. 8). This finding demonstrates two things: first, a greater number of loci than were detected by a comparison of two human individuals are polymorphic and segregate amongst human populations; and second, if the read mapping and variant calling were perfect, variation data combined with variant sequence reconstruction could be used for *de novo* computational detection of template switch mutations. Under the same constraints, the approach could also be applied to resequencing data from trios.

## Conclusions

Our generalized template switch model can explain a large number of complex mutation patterns—clusters of apparent base substitutions and indels—with a single mutation event. Although only confidently explaining 0.3% of base differences and 0.2% of indels in clusters in the human-chimp comparison, this is nevertheless a large number of individual events and far exceeds the numbers previously found in higher organisms. Note that our inferences are likely underestimates, because of our strict criteria and filtering. The model is compatible with, and significantly extends, the one previously proposed for bacteria and described replication-based mechanisms for genome rearrangements such as BIR, SRS, FoSTeS and MMBIR. Unlike previous models for short-range template switching significant pre-existing repeats or sequence similarity are not required and the process can thus create completely novel repeats (see Fig. 2; Supplementary Fig. 9). This is consistent with the reported cases of major genomic rearrangements where microidentity of only two or three bases is observed at the switch points (Lee et al., 2007; Hastings, Ira and Lupski, 2009; Costantino et al., 2013). (We prefer ‘microidentity’ to ‘microhomology’, as used by previous authors, because homology means common ancestry [Webber and Ponting, 2004] and is unnecessary for the proposed mechanisms.) We also found no evidence of the intra-strand events of the bacterial model, possibly because they would require breaking of the bonds between the leading strand template and the newly synthesized DNA. On the other hand, the most common event type that we detected, which only inverts a sequence fragment, can only be found by our generalized model.

When the template switch event does not involve loss or gain of sequence, the mutation pattern appears as a multi-nucleotide substitution (MNS). Some cases of MNSs have been explained with positive selection (Bazykin et al., 2004; Meer et al., 2010) while involvement of Pol *ζ* has been suggested to explain spatial differences in mutation frequency (Harris and Nielsen, 2014). Our results demonstrate that template switch mutations are also playing a role in the creation of clusters of adjacent substitutions. Many template switch events are associated with indels in the alignment (Supplementary Fig. 10) and the process we identified provides an alternative to the proposition of indels being mutagenic and triggering nearby base substitutions (Tian et al., 2008).

The proposed four-point model has consequences for our understanding of genome evolution and the methods used for studying it. While template switching is known to have a role in genomic rearrangements (Gu et al., 2008; Hastings, Lupski, Rosenberg and Ira, 2009; Costantino et al., 2013; Carvalho and Lupski, 2016), our analyses demonstrate that it can also take place in a local context. As such, it provides a one-step mechanism for the generation of hair-pin loops and, in combination with other mutations, provides a pathway to more complex secondary structures (Ding et al., 2014; Rouskin et al., 2014; Wan et al., 2014). The model also provides a mechanism for the evolution of existing DNA secondary structures and provides an explanation for the long-standing dilemma of exceptionally high rates for compensatory substitutions (Dixon and Hillis, 1993; Tillier and Collins, 1998; Meer et al., 2010). Interestingly, the mechanism may also maintain apparent DNA secondary structures without selective force. A number of human disease mutations (Chen et al., 2005*c*,*a*,*b*; Lee et al., 2007; Hastings, Ira and Lupski, 2009; Zhang et al., 2009) have been attributed to events that can be described by our template switch model. We also note that key mutations implicated in the *de novo* origin of the putative human protein coding gene *DNAH10OS* (Ensembl ID ENSG00000204626; Knowles and McLysaght, 2009: Fig. 4, enabling 10-bp insertion CCTCATTCCT and G*→*A substitution 2 bp downstream; see also Xie et al., 2012) can be explained by the four-point model.

A probable reason why template switch mutations have not received greater attention may be bias in commonly used analysis methods. Tight clusters of differences, the typical signature of the process, make read mapping and subsequent variant calling challenging. This is demonstrated by phase 3 of the 1kG Project (1000 Genomes Project Consortium et al., 2015), which provides significant improvements in comparison to earlier releases but, as we have shown, still contains errors and inconsistencies around the regions we have studied. High quality sequence and assembly has been vital to improving understanding of structural variation of genomes (Pendleton et al., 2015; Sudmant et al., 2015). We have shown that improving genome assemblies to the level of individual bases and short indels relative to reference sequences is needed in order to permit correct interpretation of the causes of population-level differences and of the information most commonly used to study intra- and inter-species evolution. Mapping methods that simultaneously consider multiple references are beginning to become available (e.g. Schneeberger et al., 2009; Maciuca et al., 2016), and the new mutation model we propose could be modeled and considered in future analyses. With improvements in relevant algorithms and the rapidly growing number of high quality *de novo*-assembled genomes, the full extent of local template switch events can be uncovered.

## Methods

### Discovery of four-point mutations

We downloaded the Ensembl (v.71) EPO alignments (Flicek et al., 2013; Paten et al., 2008) of six primates and included all blocks containing only one human and chimp sequence, covering in total 2.648 Gb of the human sequence and 94.8% of the EPO alignment regions. Keeping only human and chimp sequences, we identified alignment regions where two or more non-identical bases (mismatches or indels) occur within a 10-base window. For each such mutation cluster, we considered the surrounding sequence (for human and chimp, respectively, 100 and 200 bases up- and downstream from the cluster boundaries), and in accordance with our four-point model attempted to reconstruct the human query from the chimp reference with imperfect copying (allowing for mismatches and indels) of the forward strand and two freely placed template switch events. Candidate switch events were required to have high sequence similarity without alignment gaps and within the ②→③ fragment only mismatches were allowed. If exact positions of switch events could not be determined (Supplementary Fig. 11), our approach maximized the length of ②→③ fragment and reported this upper limit of the strand-switch event length. For comparison, we reconstructed the human query from the chimp reference with imperfect copying of the forward strand only (i.e. linear alignment) using the same scoring. A custom dynamic programming algorithm to determine the optimal four-point model explanation for each mutation cluster is described in Supplementary Fig. 2 and Supplementary Algorithm 1. The computational tool used for the analyses is available at http://loytynojalab.biocenter.helsinki.fi/software/fpa.

### Filtering of events

For each mutation cluster, we recorded the coordinates of the inferred template switch events and computed similarity measures for the different parts of the template switch and forward alignments as well as the differences in the inferred numbers of mutations between the two solutions; we also recorded whether the regions include repeatmasked (Smit et al., 2013–2015) or dustmasked (Morgulis et al., 2006) sites, as well as the number of different bases included in the ②→③ fragments. We then selected a set of events as high-confidence candidates using the following criteria: (*i*) the switch points ① and ④ are at most 30 bases up- and downstream, respectively, from the cluster boundaries; (*ii*) the ②→③ fragment is at least 10 bases long; (*iii*) the ②→③ fragment as well as 40-base flanking regions up-and downstream show at least 95% identity between the sequences; (*iv*) the forward alignment indicates at least two differences (of which at least one a mismatch) more than the template switch alignment (which may also contain up to 5% mismatches); (*v*) the ②→③ fragment is not repeatmasked or dustmasked and contains all four bases. As a control to assist in assessing the occurrence of false positives, we repeated the analysis without complementing the ②→③ fragment: no biological function is known for reverse repeats and we consider them a proxy for the probability of observing a repeat of particular length by chance.

### Identification of polymorphic mutations

The GRCh37 human reference and Venter Caucasian male genome sequences were aligned using LASTZ (Harris, 2007) and following the UCSC analysis pipeline (Kent et al., 2002). The four-point mutation events were identified using the same approach as with human-chimp data. The 1kG variation data from ftp://ftp.1000genomes.ebi.ac.uk/vol1/ftp/release/20130502/ were analysed using bcftools (Li, 2011) and selected regions of resequencing data from ftp://ftp.1000genomes.ebi.ac.uk/vol1/ftp/phase3/data/NA12878 were visualized using samtools (Li et al., 2009). Mutation clusters with uniform allele frequencies were identified as follows: (i) 1kG variant calls were extracted for the mutation cluster plus 10 bases of flanking region; (ii) for each locus, runs of adjacent positions with less 10% difference in global allele frequency (AF) were recorded; and (*iii*) the runs of selected length (e.g. 3) with AF between 0.01 and 0.99 were outputted. The 1kG variant alleles were reconstructed using GATK (McKenna et al., 2010). Short-read alignment data, 1kG variant calls and primate sequence alignments for the candidate template switch event loci are available at http://loytynojalab.biocenter.helsinki.fi/software/fpa.

### Other computational analyses

DNA secondary structures were predicted with the ViennaRNA package (Lorenz et al., 2011), using the command ‘RNAfold --paramFile=dna_mathews2004.par --noconv --noGU’. The length distribution (Supplementary Fig. 3) and the allele frequencies (e.g. Fig. 3d) were visualized with R (R Core Team, 2014).

## Footnotes

**Competing interest statement** The authors declare that they have no competing financial interests.

## Acknowledgements

We thank Martin Taylor for help and comments in early stages of the study, and CSC - IT Center for Science, Finland, for computational resources.

*Author contributions:* N.G. devised the extended four-point model. A.L. implemented the method and performed the analyses. N.G. and A.L. designed the study, discussed the results and wrote the manuscript. Both authors read and approved the final manuscript.

